# Deconstruction of a memory engram reveals distinct ensembles recruited at learning

**DOI:** 10.1101/2024.12.11.627894

**Authors:** Clément Pouget, Flora Morier, Nadja Treiber, Pablo Fernández García, Nina Mazza, Run Zhang, Isaiah Reeves, Stephen Winston, Mark A. Brimble, Christina K. Kim, Gisella Vetere

## Abstract

How are associative memories formed? Which cells represent a memory, and when are they engaged? By visualizing and tagging cells based on their calcium influx with unparalleled temporal precision, we identified non-overlapping dorsal CA1 neuronal ensembles that are differentially active during associative fear memory acquisition. We dissected the acquisition experience into periods during which salient stimuli were presented or certain mouse behaviors occurred and found that cells associated with specific acquisition periods are sufficient alone to drive memory expression and contribute to fear engram formation. This study delineated the different identities of the cell ensembles active during learning, and revealed, for the first time, which ones form the core engram and are essential for memory formation and recall.

## Main Text

Our cognitive system receives a continuous influx of information and processes this data to integrate select elements into our memory repository. This process involves the activation of specific cell subsets within neuronal networks during memory acquisition and requires the orchestrated reactivation of these subsets later on for memory retrieval ^1^. Contextual fear conditioning (FC) in mice, where an association between a neutral stimulus (context, Conditioned Stimulus CS) and an aversive one (electric shock, Unconditioned Stimulus US) is established, serves as a common paradigm for studying memory formation. After fear conditioning, during subsequent presentations of the CS alone, a conditioned response (CR, freezing behavior) can be observed and measured as an index of fear memory recall ^2^. Previous studies have demonstrated that targeted reactivation of cells active at the time of memory encoding across distinct brain areas can artificially induce memory recall ^3–6^, indicating the existence of the engram, the physical substrate of a memory trace ^7^. The discovery that optogenetic activation of engram cells triggers memory retrieval represented a significant advancement in our understanding of memory formation. However, what remains unanswered is why certain cells are recruited into the engram and what type of information these cells encode at the time of memory acquisition ^8^.

These questions are particularly relevant due to the coexistence of different neurons within brain regions, which are tuned to unique features of an experience. For instance, within the CA1 field of the hippocampus, studies have identified the presence of place cells ^9^, shock cells ^10^, aversive stimulus-tuned cells ^11^ and cells preferentially active during or outside of freezing episodes ^12^. Recent research has begun to investigate how these cellular populations are integrated into engrams, in particular those encoding spatial information present at the time of memory acquisition and recall ^13^. However, no studies to date have attempted to characterize if certain cell populations active at distinct periods of encoding play a disproportionate role in memory recall. Exploring this relationship represents a critical step towards understanding how the brain converts specific information from the time of encoding into memories.

A significant challenge to addressing this stems from the lack of tools capable of isolating select cells from the broader population active during memory acquisition. Specifically, prior approaches have tagged cells expressing immediate early genes (IEGs), but only over a broad temporal window which can range from several minutes to hours ^14^.

Our present work addresses this by employing state-of-the-art technology, f-FLiCRE ^15^, a novel optogenetic tool that employs light and high intracellular calcium for dual-condition tagging. This faster f-hLOV1 variant of FLiCRE, capable of unprecedented temporal resolution, allows us for the first time to tag cells active at specific moments of FC, e.g. during shock presentation or freezing bouts, and manipulate them in subsequent sessions.

We hypothesize that cells active at crucial periods of memory encoding will be preferentially recruited to the related memory engram.

### Opto-activation of FC-active dCA1 cells tagged using FLiCRE is sufficient for fear memory recall

Initially, we validated f-FLiCRE efficacy by reproducing a classic gain-of-function experiment to reveal hippocampal engrams in FC ^3,4,6^. We f-FLiCRE-tagged dorsal CA1 (dCA1) cells (Fig.1A-C) active during FC in context A (ctxA, Fig.S1A) for the 5 minutes following the first shock onset. The following day, bReaChES stimulation (in FC-tagged mice) of the tagged cells in a neutral context (ctxC) triggered freezing, indicating that the activation of FC cells supports aversive memory recall (ctxC, Fig.1D-F, Fig.S1A-E). This increase in freezing was absent in the two control groups that received the same light stimulation in ctxC but without previous tagging (NO tag), or tagging for 5 minutes in a different context in the absence of shocks (ctxB tag), highlighting the experience-dependent and context-specific nature of our results. Compared to the NO tag group, the FC-tag and ctxB-tag groups displayed a significantly higher number of tagged cells (Fig.1G-I). The FC-tagged cells were also significantly more numerous than ctxB-tagged cells, in line with the observed increase in dCA1 activity associated with high saliency events ^16,17^. Additionally, the efficacy of f-FLiCRE tagging of dCA1 active cells was validated by quantifying the tagged to infected cells ratio during kainic acid-induced seizures (67%±5%), known to massively activate neurons. Overall, these results demonstrate that engram tagging based on neuronal calcium influx reproduces previous gain-of-function findings from engram studies using IEGs-tagged cells, and supports f-FLiCRE’s potential as a novel, reliable tool for engram-tagging with considerably higher temporal resolution.

**Fig.1.**
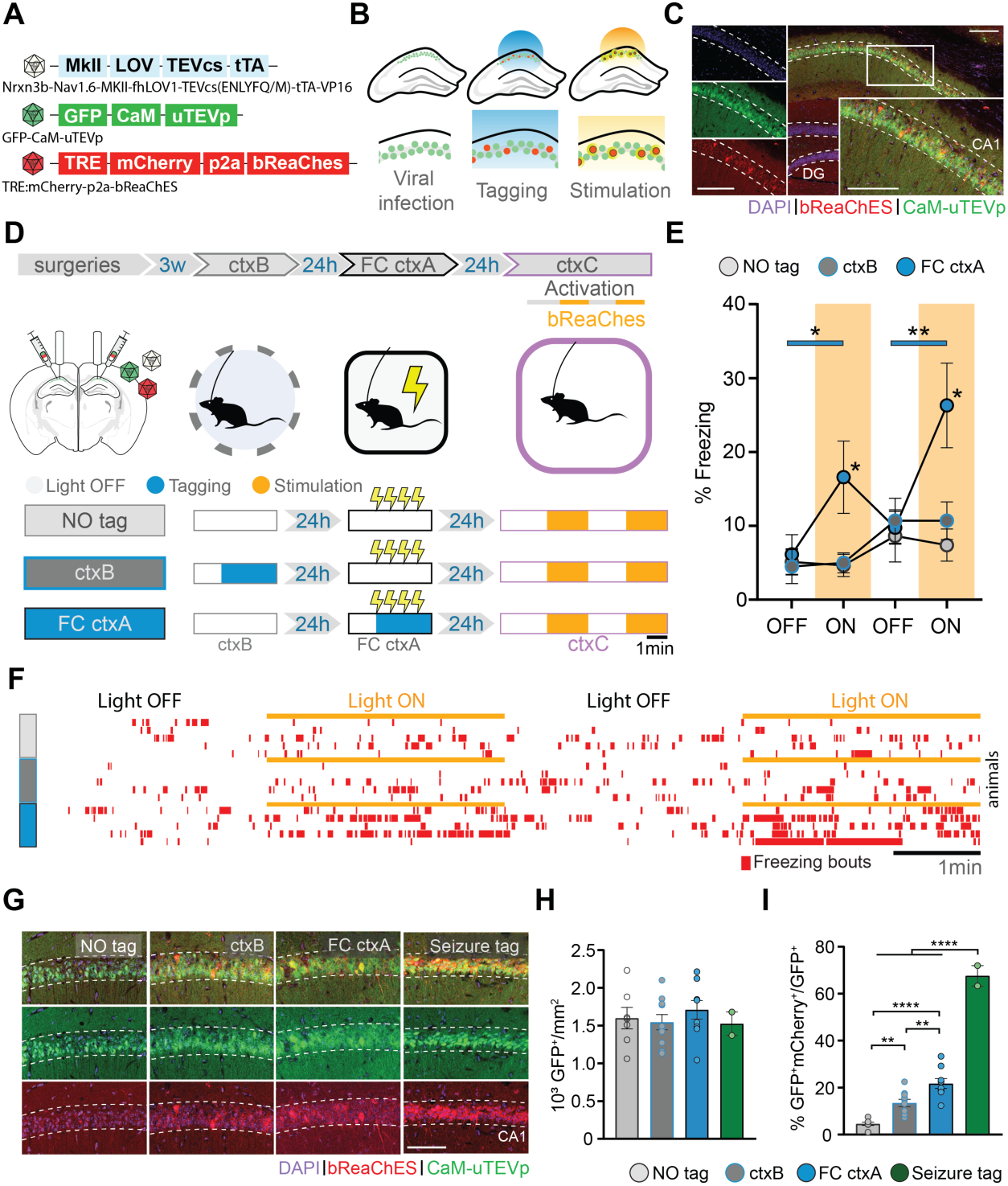
Optogenetic stimulation of f-FLiCRE-tagged dCA1 cells active during fear conditioning triggers memory recall. (**A**) f-FLiCRE viral constructs with the excitatory opsin bReaChES are injected bilaterally in the dCA1. (**B**) Cells expressing f-FLiCRE (green) are tagged using blue light. Tagged cells (red) express mCherry and bReaChES. Later stimulation with yellow light activates previously tagged bReaChes cells. (**C**) Example histology of f-FLiCRE-bReaChES infected (GFP+, green) and tagged (mCherry+, red) cells (DAPI+, blue) in the dorsal hippocampus and magnification in CA1. Scale bar: 100µm. (**D**) Schematics of the experimental protocol (top). Tagging and reactivation timelines for the three groups: “NO tag”, “ctxB”, and “FC ctxA” (bottom). (**E**) Comparison of freezing levels between groups in ctxC during light-OFF (no shading) and light-ON epochs (yellow shading). (**F**) Freezing timelines of representative individual animals. (**G**) Representative images of infection (green) and tagging (red) in the three previously described groups, as well as of a “seizure tag” control. Scale bar: 100 µm. (**H**) Density of GFP+ (i.e. infected) neurons in the dCA1 layer. (**I**) Percentage of tagged cells (GFP+mCherry+/GFP+). Each data point corresponds to the mean value for each individual animal while bars represent mean ± SEM across animals. Statistical tests are ordinary one-way or RM two-way ANOVAs depending on the case. Statistical differences are depicted with asterisks, with color-coded lines used to show between-epoch comparisons, for consecutive periods only (non-consecutive significance is not shown). *p<0.05, **p<0.01, ****p<0.001.

### dCA1 engram dissection revealed sub-engrams sufficient to trigger memory recall

To investigate whether cells preferentially active at specific times during memory encoding incorporate different information into the memory engram, we leveraged the full temporal resolution offered by f-FLiCRE. We divided FC into four tagging periods: pre-shock, shock, freezing, and no freezing (Fig.2A-C, Fig.S1F-H). Since this is the first time f-FLiCRE is used at such temporal resolution, to ensure that f-FLiCRE could tag cells active during light-ON periods, but did not tag cells active during light-OFF periods, we validate its use by repeatedly delivered bouts of 5 seconds light-ON, 5 seconds light-OFF to cultured hippocampal neurons, matching the median duration of light-ON or light-OFF periods of in-vivo “freezing” and “no-freezing” experiments (Fig.S2A). Neurons either received no stimulation (light only), 20 Hz electric field stimulation during the light-OFF periods (alternating), or 20 Hz electric field stimulation during the light-ON periods (simultaneous; Fig.S2B). While “light only” and “alternating” groups showed similar levels of tagging (TRE-mCherry reporter labelled cells), we observed a significant increase in tagged cells only in the “simultaneous” condition (Fig.S2C-E). GCaMP6f imaging in separate neurons confirmed that the elevated intracellular calcium levels returned to baseline levels in between bouts of electric field stimulation (Fig.S2F-H). These data prove that f-FLiCRE has sufficient temporal resolution and sensitivity to tag even brief bouts of neuronal activity, such as those during “freezing” and “no-freezing”.

**Fig.2.**
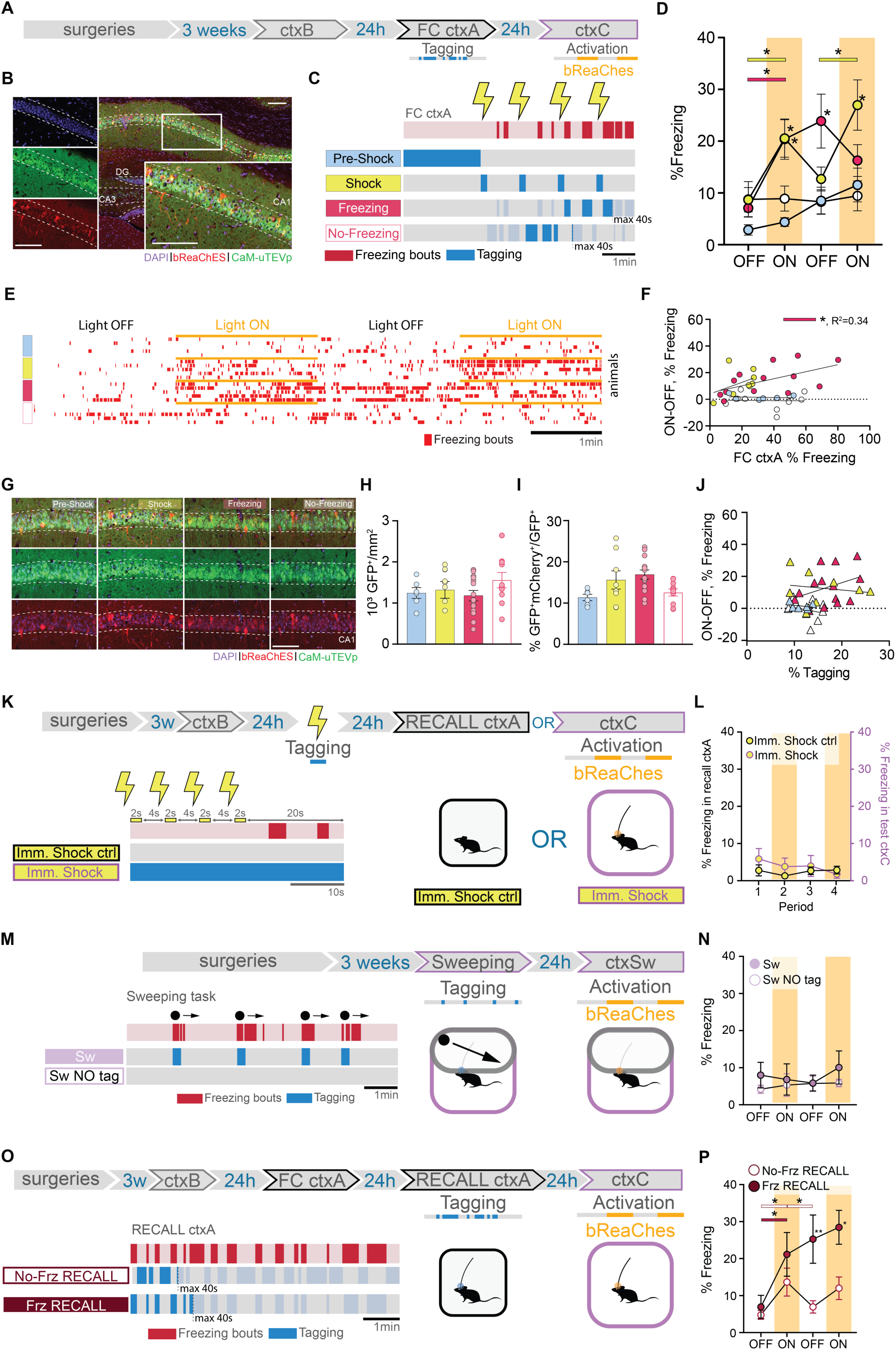
dCA1 cells differentially integrate into the engram depending on when they are active during FC. (**A**) Schematic of the experimental protocol: f-FLiCRE viral constructs with the excitatory opsin bReaChES are injected bilaterally into the dCA1; opto-tagging occurs during FC ctxA and opto-reactivation 24 hours later during ctxC. (**B**) Example histology of f-FLiCRE-bReaChES infected (GFP+, green) and tagged (mCherry+, red) cells (DAPI+, blue) in the dorsal hippocampus and magnification in CA1. Scale bar: 100µm. (**C**) Tagging strategy for “pre-shock”, “shock”, “freezing”, and “no-freezing” groups. (**D**) Comparison of freezing levels between groups in ctxC during light-OFF (no shading) and light-ON epochs (yellow shading). (**E**) Example individual freezing traces from the four different tagging groups. (**F**) Correlation between Δfreezing in ctxC and overall freezing in FC ctxA. (**G**) Representative images of infection (green) and tagging (red) in the four groups. Scale bar: 100 µm. (**H**) Density of GFP+ (i.e., infected) neurons in the dCA1 layer. (**I**) Percentage of tagged cells (GFP+mCherry+/ GFP+). (**J**) Correlation between Δfreezing in ctxC and percentage of tagged cells. (**K**) Schematics of immediate shock protocols: opto-tagging occurs during presentation of immediate shocks; “Imm. Shock ctrl” animals are tested in a recall ctxA session, while “Imm. Shock” animals are opto-reactivated in ctxC. (**L**) Freezing level comparison for the two immediate shock groups. Left axis is freezing in recall ctxA for “Imm. Shock ctrl”, while the right axis is freezing in test ctxC for “Imm. Shock” animals. (**M**) Schematics of “Sw” and “Sw NO tag’’ protocol: opto-tagging occurs during sweeping episodes in a modified ctxC and opto-reactivation 24 hours later in the same context. (**N**) Freezing level comparison for the two sweeping groups. (**O**) Schematics of “No-Freez RECALL” and “Freez RECALL” groups: opto-tagging occurs during RECALL ctxA 24 hours after FC ctxA and opto-reactivation occurs during ctxC, another 24 hours later. (**P**) Freezing level comparison for the recall groups. Each data point corresponds to the mean value for each individual animal while bars represent mean ± SEM across animals. Statistical tests are ordinary one-way or RM two-way ANOVAs depending on the case. Statistical differences are depicted with asterisks, with color-coded lines used to show between-epoch comparisons, for consecutive periods only (non-consecutive significance is not shown). *p<0.05, **p<0.01.

The four subdivisions (“Pre-Shock”, “Shock”, “Freezing”, and “No-Freezing”) correspond to crucial periods of FC that together form the experience. During the pre-shock period, mice explore a novel environment and the dCA1 cells process contextual information, prior to aversive stimulus exposure. Shock delivery constitutes the US which triggers a fearful association with the context. Following shock delivery, mice engage in different behaviors, including freezing. We measured bouts of freezing as a behavioral readout for the internal fearful state of the animal, which we hypothesized influences memory acquisition. Hence, we tagged cells active during freezing, or outside of freezing bouts using a closed-loop system that relies on DeepLabStream for online animal pose estimation and intermittent light delivery (^18^, Fig.S1F-H). This choice is further supported by the fact that shock-responding as well as freezing-activated cells have been previously described in the hippocampus ^10,12^. To ensure distinct tagging between the “shock” and “freezing” conditions, we controlled that “shock-tagging” epochs included minimal freezing behaviour (Fig.S1G).

We assigned different batches of animals to the four mentioned groups and reactivated f-FLiCRE-tagged cells in a neutral context (ctxC) one day after (Fig.2A-C). The optogenetic reactivation revealed that only "shock" or "freezing" tagged mice exhibited elevated freezing behavior during light-ON epochs, while “pre-shock” or “no-freezing” tagged mice did not (Fig.2D-E, supplementary movies). Apart from freezing, the different groups exhibited similar behaviors in ctxC (Fig.S3A), suggesting that optogenetic stimulation specifically triggers fear memory-like responses. This result suggests that cell populations active at certain distinct encoding periods (“shock” and “freezing”) play an important role in memory recall, while others do not (“pre-shock” and “no-freezing”).

The number of infected cells and ratio of tagged to infected cells was comparable across all groups (Fig.2G-I), and ctxC test Δfreezing was not influenced by the tagged-to-infected ratio in any condition (Fig.2J). Because some individual animals from the “shock”-tagged and “freezing”-tagged groups displayed high tagged/infected ratios, we confirmed that their exclusion did not impact the behavioral differences observed (Fig.S3B-E), further demonstrating the importance of content over quantity of tagged cells.

While “shock” and “freezing” cells can both drive freezing behavior if reactivated, we found significant differences between them. First, FC “freezing”-tagged mice did not return to pre-activation freezing levels during the second light-OFF period, maintaining fear memory recall in absence of further opto-stimulation, while “shock”-tagged mice did. Second, the correlation between the percentage of freezing during FC and the light-driven increase in freezing during test in ctxC (Δfreezing = %freezing ON-OFF) was significant only in the “freezing”-tagged group, and not the “shock”-tagged or any other tagged group (Fig.2F, Fig.S3F-H). This correlation reproduces a natural feature of fear memory recall (Fig.S3I) and suggests that the reactivation of “freezing”-tagged cells may engage unique dynamics involved in fear memory recall that “shock”-tagged cells cannot.

Our “pre-shock” results are perhaps surprising considering that the pre-shock period is essential to creating the US association with the context ^19^. To eliminate the possibility that contextual information is not sufficiently processed at the time of tagging within these cells, we performed an additional experiment tagging “pre-shock” cells only after an 8-minute long pre-exposure period, and obtained similar results (Fig.S4A-C). This experiment confirmed that cells active during FC before US delivery are unable to trigger memory recall and may be explained by US induced-remapping of dCA1 neuronal representations ^20–22^.

To ensure that the behaviors triggered by the “shock” and “freezing” groups are specifically related to memory, we conducted two additional experiments.

In the first experiment, we tagged “shock”-responding cells using an immediate-shock protocol, in which the shock was delivered immediately after placing the mice in ctxA. Under these conditions, mice are unable to form a fear association with the context ^19^, as shown by the absence of freezing behavior when re-exposed to the same context the following day (Fig.2K-L, “Imm. Shock ctrl”). Reactivating “shock”-responding cells tagged during immediate-shock protocol did not elicit fear behavior in ctxC the next day, confirming that “shock”-responding cells must be linked to an associative fear memory to drive a freezing response. In contrast, and supporting the results shown for the main “shock” group (Fig.2D), when the same delay between shocks was applied after 2 minutes of context exposure, mice successfully formed a memory association (Fig.S4D-E), and reactivating these cells induced freezing behavior in a neutral context (Fig.S4F-G).

In the second experiment, we tagged “freezing” cells in a sweeping task (Fig.S1A, Fig.S4H-I) (25). Mice freeze as an instinctive response to a perceived overhead threat without an associative memory being formed with the context, as evidenced by the absence of freezing when mice are returned to the same context the next day (Fig.2M-N, “Sw NO tag” control group). Reactivation of these cells in the same context one day later did not elicit freezing behavior (“Sw” group), suggesting that the effect observed in the “freezing” group in the FC task is memory-related (Fig.2D).

As the majority of ^3,4,6,23,24^, but not all (see also ^25^) studies targeting the engram have focused on tagging cells during memory acquisition, little is known about the composition of engram cells during memory recall.

To investigate whether specific cells are preferentially involved in memory recall the day after memory formation, we tagged cells active during freezing (“Frz RECALL” group) or no-freezing (“No-Frz RECALL” group) periods as done previously, but this time during a memory recall session in ctxA (Fig.2O). Opto-activation of both groups in a neutral context the following day resulted in increased freezing behavior (Fig.2P). This suggests that, unlike during memory formation, during the memory test, engram cells can be active both during and outside of freezing bouts. This leads to the conclusion that, while only certain acquisition cells are selected to become part of the memory engram, engram cells can be activated at any point during memory recall, whether memory-related behaviors are being expressed or not.

Interestingly, similarly to the FC-”freezing”-tagged group, recall-”freezing”-tagged mice did not return to control freezing levels during the second OFF period, further supporting the idea that different sub-engrams contribute to the engram in different ways.

Within these additional groups (“long-exp.”, “Pre-shock”, “no-freez recall”, “freez-recall”, “sweeping”, “Imm. Shock”, and “Imm. Shock ctrl”), some showed differences in expression and tagging (Fig.S4J-K), there was no effect of % tagging on test Δfreezing in any condition (Fig.S4L-O).

### dCA1 engram dissection revealed that the same sub-engrams are also necessary for memory recall

To further demonstrate the distinct integration into the engram of the four subpopulations of cells identified during memory acquisition, we conducted a loss-of-function experiment. We replaced bReaChES with eNpHR3.0, an inhibitory opsin, within the f-FLiCRE system (Fig.3A). Instead of using the ctxC test session, we introduced a recall session in ctxA, during which we could intermittently inhibit tagged cells. We tagged cells activated during “pre-shock”, “shock”, “freezing”, or “no-freezing” conditions, replicating the same experimental setup as previously described, and included a “NO tag” control group to assess the efficacy of the new reporter protein (Fig.3B-C). Our findings revealed that only inhibition of the “shock” or “freezing” tagged cells, but not the “pre-shock”, “no-freezing” nor “NO tag” cells, led to a significant decrease in freezing levels, and that this effect persisted during the second OFF period in both groups (Fig.3D). The percentage of infected cells was comparable between groups, percentage tagged cells was similar for all groups except for the NO tag group (Fig.S5A-C), and did not correlate with Δfreezing for any group (Fig.S5D). These results indicate that the “shock” and “freezing” (but not “pre-shock” and “no-freezing”) subsets of engram cells are not only sufficient (Fig.2), but also necessary for the behavioral expression of fear memory. Furthermore, paralleling the excitatory results (Fig.2F), freezing in FC ctxA correlated with Δfreezing only in the “freezing” group (Fig.S5E).

**Fig.3.**
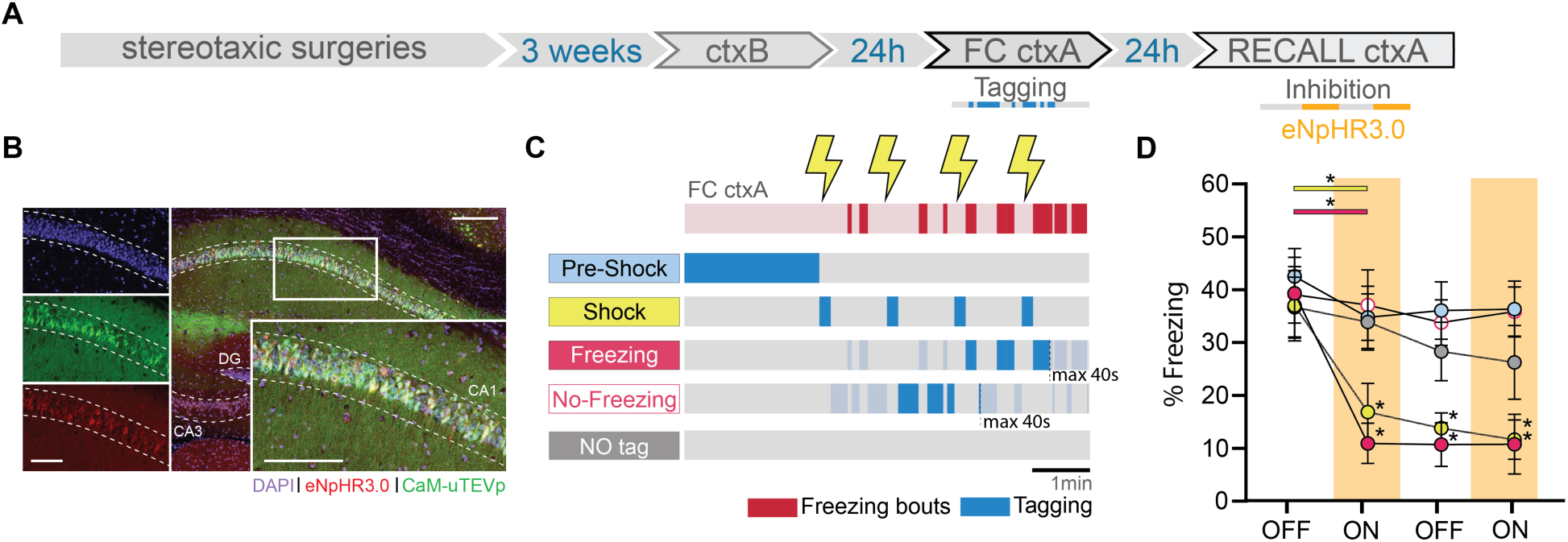
dCA1 cells whose reactivation is sufficient for memory recall are also necessary for memory recall. (**A**) Schematic of the experimental protocol: f-FLiCRE viral constructs with inhibitory opsin eNpHR3.0 are injected bilaterally in mouse dCA1; opto-tagging occurs during FC ctxA and opto-inhibition during recall ctxA 24 hours later. (**B**) Example histology of f-FLiCRE-eNpHR3.0 infected (GFP+, green channel) and tagged (mCherry+, red channel) cells (DAPI+, blue channel) in the dorsal hippocampus and magnification in CA1. Scale bar: 100µm. (**C**) Schematic of the tagging protocols for “pre-shock”, “shock”, “freezing”, “no-freezing” and “NO tag” inhibition. (**D**) Comparison of freezing levels between groups in recall ctxA during light-OFF (no shading) and light-ON epochs (inhibition, red shading). RM two-way ANOVA. Statistical differences are depicted with asterisks, with color-coded lines used to show between-epoch comparisons, for consecutive periods only (non-consecutive significance is not shown). *p<0.05.

### Calcium imaging revealed the presence of non overlapping dCA1 neuronal ensembles

To understand the relationship between dCA1 “sub-engrams” active at the time of encoding and their activity in memory recall, in a separate experiment, we characterized single neuron activity by recording dCA1 calcium dynamics in freely moving animals during ctxB, FC ctxA, recall in ctxA, and in ctxC (Fig.4A-B, Fig.S6A). We assigned the cells detected to groups according to their activity in FC ctxA. We tailored our method to best match f-FLiCRE tagging, as we aimed to study the activity of the same cells that would have been tagged in our optogenetic experiments (Fig.4C). We first analyzed the overlaps between pairs of subpopulations (i.e. the proportion of cells classified as part of both). At encoding, every pair of f-FLiCRE-like cell groups showed significantly less overlap than chance, except for the shock group, which showed chance-level overlap with the freezing and non-freezing groups (Fig.4D). This indicates that the 4 populations are largely separate from one another, despite the greater observed topographic proximity between “shock” and “freezing” cells relative to the other groups (Fig.S6B-D). In a non-shocked context (ctxB), the overlap was non-significant between cells active in the first 2 minutes or later on (Fig.S6E-F), demonstrating that the significant non-overlap between “pre-shock” and other populations was not due to FC-independent intra-experiment changes in activity. We also analyzed f-FLiCRE-like groups of cells active at recall during freezing or no-freezing (“Frz RECALL” and “No Frz RECALL” cells) and found that these two subpopulations showed higher than chance overlap levels (Fig.4C-D).

**Fig.4.**
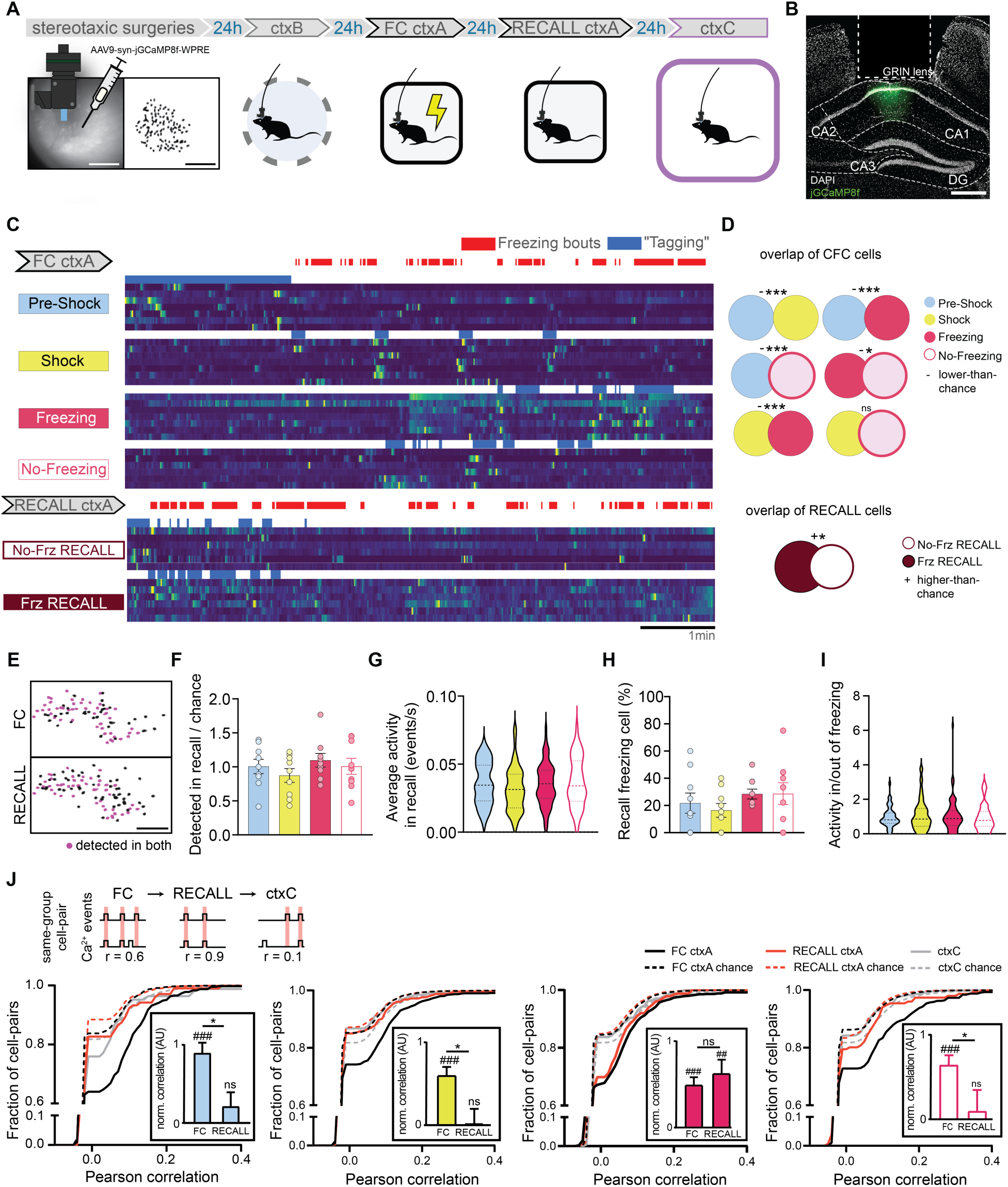
In-vivo calcium imaging of FC ctxA tagged cells. (**A**) Schematic of the experimental protocol: jGCaMP8f is injected into dCA1, a relay lens is placed on top of it and a baseplate is installed to allow imaging using the miniscope. Five weeks later, dCA1 cells are imaged across three experiments. (**B**) Example histology of injection and lens placement. DAPI is in white, jGCaMP8f in green. Scale bar: 400 µm. (**C**) Example calcium traces from the four different FC ctxA f-FLiCRE-like cell groups, as well as the two RECALL ctxA ones. Blue represents the periods used to determine which cells belong to the corresponding group, and would have been tagged using f-FLiCRE. (**D**) Overlap between cell groups from the same experiment, top: FC, and bottom: RECALL. ‘+’ denotes higher-than-chance overlaps, ‘-’ lower than chance overlaps, ‘n.s.’ denotes no significant difference from chance. (**E**) Example cell tracking between two experiments using CellReg. Scale bar: 100 µm. (**F**) Number of cells per animal from FC ctxA cell groups detected in RECALL ctxA over chance number by cell group. (**G**) Average calcium events per second in RECALL ctxA for tracked cells by cell group. (**H**) Percentage of FC ctxA groups cells that are freezing cells in Test ctxA. (**I**) Activity ratios per animal of FC ctxA groups for cells inside/outside of RECALL ctxA freezing bouts. (**J**) Cumulative distributions of same-group cell-pair Pearson correlations in FC ctxA (black lines), RECALL ctxA (red lines) and ctxC (gray lines). Chance-level correlations are shown with dotted lines. Statistical tests are comparisons to bootstrapped distributions for overlaps, and ordinary one-way ANOVA otherwise. *p<0.05, ***p<0.001, ##p<0.01, ###p<0.001.

We then tracked the activity of the four subpopulations detected in FC ctxA during RECALL ctxA and ctxC (Fig.4E), hypothesizing that “shock” and “freezing” cells would be preferentially reactivated. However, in RECALL ctxA, we found no differences between any of the f-FLiCRE-like groups in terms of proportion of tracked cells, overall activity, activity during freezing bouts, or proportion that were also “Frz RECALL” cells (Fig.4F-I). These analyses show that neuronal reactivation during memory recall is uniform across the groups and suggest that dCA1 engram reactivation relies on more complex neuronal dynamics. To reveal these dynamics, we computed the correlations in calcium events during FC ctxA, RECALL ctxA, and ctxC between same-group cell pairs (Fig.4J) and found that, for all groups, same-group cell pairs were highly and significantly correlated during FC ctxA. However, only “freezing” cell pairs maintained a significant correlation in RECALL ctxA, while others did not. In ctxC correlations were non-significant for all groups (Fig.4J). To verify the validity of these results, we ran all the previous analyses again, but switched the method for defining neurons’ identity to a statistical criterion (Fig.S7A, see Methods) instead of an activity-based one, and obtained identical or highly similar results (Fig.S7B-G).

Overall, these results suggest that, although several distinct ensembles of neurons emerge during memory formation, only the “freezing” cells are reactivated as an ensemble during memory recall.

## Conclusions

This set of experiments showed for the first time that specific external stimuli (i.e. shock delivery, US) and internal states (i.e. freezing behavior, CR), which occur at specific moments during memory acquisition, are associated with distinct, non-overlapping neuronal ensembles that are both necessary and sufficient for fear memory recall. In contrast, cells active before these external stimuli are presented (i.e. before the shock delivery) or outside of these internal states (i.e. during non-freezing behaviour) do not drive observable signs of fear memory recall in mouse behavior.

Recent technological advancements, including FLiCRE technology (but see also references for Cal-Light and FLARE ^26–29^), have paved the way for differentiating between and tagging cell populations with higher temporal resolution. This was unachievable with drug- and IEG-based engram tagging, considering that both the timing of drug delivery and that of IEG-derived protein production occur at a larger time scale than the acquisition window and that the expression of IEGs changes across brain regions, tasks, and moments ^8,30,31^. Instead, FLiCRE tagging relies on light delivery- and calcium levels-dependency. The new f-FLiCRE variant that we validate and use here presents even faster kinetics (^15^, Fig.S2), allowing us to target, for the first time, cells active in short, intermittent moments (e.g. “freezing” bouts) that require the ability to rapidly switch tagging on and off.

The hippocampus, particularly its CA1, CA3, and DG areas, has been central to engram research ^3,4,13,23,24,26^. In our study, we focused on the hippocampal CA1 field due to its known involvement in both formation and recall of recent contextual fear memories ^32^. While CA1 is believed to handle the contextual aspect of the experience, it also conveys valence information. Cells such as place cells, shock cells, and freezing cells have been observed in the CA1 field, prompting inquiry into their potential differential engagement in the engram ^9–12^.

In this study we found that active cells tagged during the “pre-shock” period, when mice are presumably processing the contextual component of the experience, failed to trigger freezing behavior the day after the opto-stimulation, in both short (Fig.2D) or long-exposure (Fig.S4A-C) context protocols. These results likely stem from experience-driven changes in neuronal activity, previously reported as remapping of place cells following the introduction of a fearful stimulus into a context ^20,33^, engagement of distinct ensembles to record places and experiences ^13^ or drift in neuronal activity ^34,35^. Confirming this hypothesis, calcium imaging showed that “pre-shock” cells are significantly less active after shock and form a subpopulation distinct from and non overlapping with the other groups. This shift in recruited neurons immediately following shock delivery has also been previously observed in the Prelimbic cortex ^36^, where f-FLiCRE inactivation of “post-shock” cells, but not “pre-shock”, impaired memory recall the day after. However, we also showed that even after shock-driven neuronal reconfiguration, not all cells active (see “no-freezing” group) will be selected for the engram. Reports of artificial memory recall driven by reactivation of hippocampal cells active minutes or hours before FC ^27^ are not contradictory with the failure to recall we observed when reactivating “pre-shock” or “no-freezing” neurons. In fact, putative preallocated neurons could be active at any moment during training, including any of the four periods we used for tagging.

We then showed that both “shock” and “freezing” cells are necessary and sufficient for fear memory recall, leading to the conclusion that, during memory encoding, these two populations are selected to become part of the engram. We observed a topographical proximity between the two, that (Fig.S6) could reflect their sharing of similar inputs or outputs, an intriguing potential explanation for their recruitment during training ^37,38^. Despite that, we also showed that differences exist between the two: 1) that only the opto-reactivation of dCA1 “freezing” cells and not “shock” cells led to sustained fear memory retrieval even after optogenetic stimulation had ended (Fig.2D); 2) that artificial recall in the “freezing” group correlated with fear levels at encoding (Fig.2F, Fig.S5E), a typical characteristic of naturally recalled memories (Fig.S3E); 3) that “shock” and “freezing” cells formed highly distinct, non-overlapping populations (although spatially close) (Fig.4D). These differences led us to hypothesize that, although both populations are selected in the engram (i.e. they are both necessary and sufficient to recall the fear memory), only dCA1 “freezing” cells, and not “shock” cells, engage in unique memory signature dynamics. To investigate this, we computed the correlations of calcium traces between cells from the same groups across days. We did indeed observe a strong correlation between “freezing” cells, which persisted during fear recall, suggesting that “freezing” cells, unlike “shock” cells, were being reactivated as ensembles ^39,40^.

While optogenetic reactivation of certain neuronal populations can trigger memory recall, our calcium imaging analysis revealed that neither population is significantly more active than others during the memory recall under normal conditions. This finding underscores the complexity of neuronal ensemble response patterns, which may rely on dynamics not revealed by single-cell analysis ^39,41^. It also highlights the need for a deeper understanding of the dynamics that emerge during prolonged ensemble optogenetic stimulation, especially in light of previous evidence that optogenetically tagged engrams can, through reactivation, eventually reconfigure to adopt endogenous firing patterns ^42^.

During FC training we identified 4 distinct and non overlapping (average ∼1%) populations of cells. Out of them, only two (“shock” and “freezing”) were capable of inducing memory recall upon reactivation. In contrast, RECALL cell populations showed greater overlap (∼15%), with all tagged cells being capable of triggering artificial memory recall, showing that a more homogeneous population is engaged at memory recall.

Altogether our results reveal that engrams are formed by selecting specific subpopulations of cells active at specific moments of acquisition (i.e., “shocks” and “freezing”). These non-overlapping populations present distinct recall dynamics that are only revealed at higher levels of analysis.

The traditional conceptualization of the engram is evolving, and a deeper understanding of the complex interplay between different neuronal populations is needed to fully unravel the mysteries of memory. Our findings revealed the neuronal components of the fear memory engram and open new doors for understanding memory engrams and the complex dynamics underlying memory formation and recall.

## Acknowledgments

We thank Colleen J. Gillon for the helpful discussions, comments, and revisions. We also would like to thank Gabrielle Girardeau, Bianca Silva and Victor Cazares for commenting on a previous version of the paper.

## Author contributions

CP and GV designed the experiments. CP, FM and NM conducted all optogenetics experiments. CP conducted all calcium imaging experiments and subsequent analysis. CP, FM, NT, PF and NM conducted all histology and subsequent analysis. RZ and CKK performed f-hLOV1 f-FLiCRE experiments in neuron culture. MB, IR and SW prepared the viruses for all f-FLiCRE experiments. GV and CP wrote the paper.

## Data and materials availability

All data are available in the main text or the supplementary materials. Codes can be found online on Zenodo (https://doi.org/10.5281/zenodo.14276916).

## Statistics

Details of statistical tests are available in supplementary table file 1.

## Materials & Methods

### Animals

All mice used were C57BL/6NRj x 129/SV first-generation hybrids, both male and female, obtained by in-house breeding of C57BL/6NRj and 129/SV (Janvier labs). Mice were housed in groups of 2 to 4 animals (including post-surgery), on a standard food and water diet (ad-libitum), and a 12h/12h light/dark cycle. Mice were 8-9 weeks old at the time of surgery (12-13 weeks old when perfused). All procedures were performed in accordance with the official European guidelines for the care and use of laboratory animals (86/609/EEC), following the Policies of the French Committee of Ethics (Decrees n° 87–848 and n° 2001–464) and after approval by ethical committee (reference: 2023-05).

### f-FLiCRE viral production

f-FLiCRE expression vectors (Addgene) were packaged into AAV serotype DJ (cell Biolabs #VPK-400-DJ) at St. Jude Children’s Research Hospital. Adenoviral helper genes were provided using the plasmid pHGTI-adeno1 ^43^. Plasmids were transfected into Adherent 293T cells (ATCC CRL-3216), 10x15 cm dishes per construct, at a 1:1:1 molar ratio using PEI-polyethyleneimine “max,” (Polysciences #24765). Media was changed to serum-free DMEM at 16 hours post-transfection. Cell supernatants and pellets were harvested 72 hours post-transfection. Cell pellets were lysed by 5 freeze-thaw cycles, lysate was collected and diluted in serum-free DMEM to a volume of 40mL and pegylated with 40% PEG8000 (Fisher) at a 1:5 overall volume (10mL PEG). Supernatants were directly pegylated with 40% PEG8000 at a 1:5 overall volume. Pegylated lysates and supernatants were incubated at 4°C for two hours and then centrifuged for 30 minutes at 4°C (4000g). The PEG-containing pellets were resuspended in 10 mM Tris 10 mM Bis-Tris-Propane (pH 9). The resuspended sample was treated with benzonase (Sigma) for 1 hour at 37°C and loaded onto a two-step (1.5g/mL:1.3g/mL) CsCl step gradient ^44^ in a thick wall ultracentrifuge tube (Beckman Cat# 360743). Samples were loaded onto an ultracentrifuge (Sw32Ti rotor) and spun at 24,600rpm at room temperature for 20 hours. The full particle containing fraction was isolated, loaded onto a dialysis cassette (Thermo cat# 66810), and dialyzed 3x in 1xPBS. The dialyzed virus was collected, filtered through a 0.2µM filter, and concentrated by a 100 kDa filter (Amicon) to a volume of 0.5mL. AAV vectors were titered by qPCR using serial dilutions of purified virus compared against linearized plasmid standard references. Viruses were aliquoted and stored at -80°C until use. All four f-FLiCRE plasmids were a gift from Alice Ting and are available on Addgene (Addgene plasmid #163031, #163032, #163037, and #158703).

### f-FLiCRE virus preparations

Two different viral preparations were used, depending on which reporter protein was needed (mCherry, and either an excitatory or inhibitory opsin). The excitatory preparation consisted of a 1:1:1 mix of AAV-DJ packaged with:

● Nrxn3b-Nav1.6-MKII-f-hLOV1-TEVcs(ENLYFQ/M)-tTA-VP16
● GFP-CaM-uTEVp
● TRE:mCherry-p2a-bReaChES

The inhibitory preparation consisted of a 1:1:1 mix of AAV-DJ packaged with:

● Nrxn3b-Nav1.6-MKII-f-hLOV1-TEVcs(ENLYFQ/M)-tTA-VP16
● GFP-CaM-uTEVp
● TRE-mCherry-p2A-eNpHR3.0-TS

### f-FLiCRE stereotactic surgeries

For all surgeries, mice were anesthetized with an i.p. injection of Ketamine/Xylazine (100/10 mg/kg) and received local anesthesia via subcutaneous injection of lurocaïne (4 mg/kg) as well as analgesia via subcutaneous injection of buprenorphine (0.1 mg/kg). After every surgery, mice were rehydrated and received analgesia in the form of subcutaneous injections of both 5% glucose water (150uL) and Meloxicam (20 mg/kg).

### f-FLiCRE viral injection

A glass micropipette pulled from a glass capillary (1.5 OD x 1.17 ID x 100 L mm, GC150T-10, Harvard Apparatus) was attached to a 10uL Hamilton microsyringe (701N, Hamilton) filled with water. Virus was loaded into the micropipette, separated from water by an air bubble. A syringe pump (Single syringe pump,

Fischerbrand) was used to control injection. Mice were injected bilaterally into the dCA1 at coordinates −2 mm anteroposterior (AP); ± 1.3 mm mediolateral (ML); –1.4 mm dorsoventral (DV). The glass micropipette was lowered before injecting 800nL of one of the two f-FLiCRE viral preparations at 0.1μL/min. We then waited 5 minutes before slowly withdrawing it. The wound was then sutured and the mice were allowed to rest.

### Optic fiber implantation

Optic fibers (200µm diameter, FP400 ERT, Thorlabs) mounted on ceramic ferrules (CFLC230, Thorlabs) were slowly lowered bilaterally to locations above the injection sites at -2 mm AP; ± 1.3 mm ML; –1.15 mm DV. Two screws were attached to the skull beforehand. Finally, dental cement (black Ortho-Jet, LANG) was applied to secure the fibers in place and close the incision.

### f-FLiCRE general experimental timeline

To allow for sufficient viral expression, experiments started 4 weeks after viral injection. Optic fiber implantation was performed on week 3 to allow both sufficient post-surgery recovery time and to limit the amount of time mice lived with the fibers implanted. All experiments were then conducted over 2, 3, or 4 days of the 4th week (depending on the group, one experiment per day). Experiments were conducted during the light phase, typically around 9 am to 12 pm. Mice were sacrificed by intracardiac perfusion in the afternoon following their last experiment.

### f-FLiCRE in-vitro experiments

Embryonic day 18 rat hippocampal neurons (Brain Bits) were prepared using the manufacturer’s recommended protocol and provided reagents. Neurons were plated on 35-mm glass bottom dishes (Cellvis) coated with 0.1 mg/ml poly-D-lysine (Millipore) dissolved in 1x borate buffer (Thermo Scientific). Dishes were rinsed with deionized sterile water and dried in the tissue culture hood. At days in vitro (DIV) 5, neurons were infected with either AAV5-hSyn-GCaMP6f (2 uL per dish; Addgene 100837-AAV5) or AAV2/1-hSyn-FLiCRE viruses (300 uL per dish; in-house crude supernatant AAV). The dishes were wrapped in aluminum foil and placed in the incubator. At DIV 14, neurons were treated under various conditions. Prior to all experiments, neurons were treated with 50 uM APV and 20 uM NBQX. Electric field stimulation was delivered using a stimulus isolator (WPI) connected to two platinum iridium wires (Alfa Aesar) shaped into rectangles that contacted opposite sides of the dish. Stimulation pulses (5-ms width) were delivered at 20Hz, using the 10-mA current setting on the stimulus isolator. Electric field stimulation was delivered at 20 Hz for 5-s on, and then 5-s off. Light delivery was performed using a 470-nm LED (Thorlabs) placed directly below the dish (5-mW output power measured at the LED). Light was delivered continuously for 5-s on, and then 5-s off. All timing was controlled using an Arduino to send time-locked TTL pulses to the stimulus isolator and LED controller box. All imaging (both timelapse and fixed) was performed using a Keyence BZ-X810 fluorescence microscope. Prior to imaging f-FLiCRE expression, neurons were fixed using ice-cold methanol. GCaMP/GFP was visualized using a 470/40-nm excitation filter and a 525/50-nm emission filter, and mCherry was visualized using a 545/25-nm excitation filter and a 605/70-nm emission filter. Images were analyzed using CellPose v2.2.3, and custom scripts in Fiji/ImageJ v2.9.0 and MATLAB vR2020b. CellPose was used to identify cellmasks from the green fluorescence channel. Cellmasks were then exported to Fiji, where the mean GFP and mean mCherry cell fluorescence were calculated for all images. The area was also calculated for all cells. MATLAB was then used to exclude cells under a certain size threshold, and to calculate the mean number of mCherry+GFP+ cells in each FOV. Prism10 was used for statistical analysis and graph making.

### Contexts for behaviors

Behaviors were conducted in four different settings: Context A (ctxA) was a 24 W x 24 L x 40 H cm glass-walled square inside a plain, white-walled, sound-proof conditioning chamber (iMETRONIC) with dim white lighting. The chamber had a metal grid floor for foot shock delivery. Context B (ctxB) was a 24 ID x 40 H cm black and white walled cylinder inside the same conditioning chamber. The chamber was fitted with a plastic, textured floor, and lit with dim blue light. Context C (ctxC) was a white-walled and floored 47 W x 47 L x 35 H cm arena with dim white lighting. For sweeping experiments, we reused the ctxC chamber, fitted a 24’’ LCD screen 40 cm above the floor. The screen displayed a plain-white background on low luminosity, to match the overall lighting intensity of ctxC. In all contexts, a camera filming at 30 fps was fitted about 50 cm above the floor to record the mice’s behavior. Since the LCD screen blocked the way in the sweeping context, the camera was offset by about 30° to be able to film inside the chamber. All chambers were wiped with 70% ethanol before animal introduction.

### Behavioral experiments

For ctxB sessions, animals were placed in ctxB for a total of 7 minutes. For FC ctxA sessions, animals were placed in ctxA for a total of 7 minutes. During this time, animals received four 0.2mA, 2s foot shocks at minutes 2, 3, 4, and 5. For fear memory recall, animals were placed in ctxA for a total of 8 minutes. For ctxC sessions, animals were placed in ctxC for a total of 8 minutes. For the sweeping task, animals were placed in the sweeping chamber for a total of 8 minutes. A preliminary study helped us determine the correct sweeping parameters to reproducibly trigger freezing bouts in animals: the dot was 2.7 cm in diameter and displayed 40 cm above the animal. The dot’s start and end points were randomized, but the dot always moved at 7° per second, traversing the screen in about 10 seconds. We wrote custom Matlab code using PsychToolbox to manually trigger sweeps of the dot across the screen. This was done at random intervals, and only when the animal was standing in the middle of the arena. Animals froze for almost the whole dot sweep. Hence, four 10-second sweeps were displayed, at varying times (sweeps were only triggered when the animal was alert, crossing the arena’s center). For the sweeping test, animals were placed back in the sweeping chamber for a total of 8 minutes. For “Imm. Shocks”, animals were placed in ctxA for a total of 40 seconds. Animals received four 0.2mA, 2s foot shocks at seconds 2, 8, 14, and 20. For “Grouped Shocks”, animals were placed in ctxA for a total of 5 minutes. Animals received four 0.2mA, 2s foot shocks at 2:00, 2:06, 2:12, and 2:18.

### Excitatory (bReaChES) f-FLiCRE laser stimulation

The optogenetic groups using the mCherry-P2A-bReaChES reporter were f-FLiCRE-tagged using a 473 nm (blue) laser shone continuously at 5mW. After 24 hours, f-FLiCRE-tagged neurons were reactivated using a 589 nm (yellow) laser shone at 5mW in 20 ms flashes at 10 Hz.

### Excitatory f-FLiCRE-tagging protocol

**“**FC ctxA” tag animals were tagged during FC, for the 5 minutes following the first shock. “ctxB” tag animals were tagged for the last 5 minutes of ctxB. “NO tag” animals did not receive any blue light. “Pre-shock”, “shock”, “freezing”, and “no-freezing” tag animals were all tagged during FC: “pre-shock” animals were tagged during the two minutes preceding the first shock. “Shock” animals received 10 seconds of blue light during and after every shock, for a total of 40 seconds of tagging. “Freezing” animals received blue light whenever they froze after the third shock, totaling 40 seconds of blue light delivery. This start time was used to prevent tagging during uncertain periods when animal position estimation was too unreliable to ensure accurate tagging of freezing behavior only. “No-freezing” animals received blue light whenever they were moving enough to not be considered freezing, starting after the second shock, for a total of 40 seconds of blue light delivery. This start time ensured the same 40 seconds of total tagging could be achieved as for the freezing and shock groups. “RECALL Frz” tag and “RECALL no-Frz” animals were tagged during the memory recall session. These animals received 40 seconds of blue light during either freezing or no-freezing bouts, respectively, without additional conditions. For “sweeping” tag animals, blue light was delivered during the four 10-second presentations of the sweeping dot, totaling 40 seconds of tagging. For “Imm. Shocks”, animals received 40 seconds of blue light for the entire duration of the FC session. For “Grouped Shocks”, animals received 40 seconds of blue light between 2:00 and 2:40.

### Excitatory f-FLiCRE manipulation protocol

Optogenetic reactivation experiments were either conducted in ctxC or the sweeping chamber (“Sw” group). In both cases, mice were placed in the chamber for a total of 8 minutes, receiving yellow light stimulation in alternating 2-minute bouts (2 minutes OFF, 2 minutes ON, 2 minutes OFF, 2 minutes ON, 20 ms pulses at 10 Hz).

### Inhibitory (eNpHR3.0) f-FLiCRE laser stimulation

The optogenetic groups using the mCherry-P2A-eNpHR3.0 reporter were f-FLiCRE-tagged using a 473 nm (blue) laser shone continuously at 5mW. After 24 hours, f-FLiCRE-tagged neurons were inhibited using a 589 nm (yellow) laser delivered continuously at 5mW.

### Inhibitory f-FLiCRE-tagging protocol

**“**Pre-shock”, “shock”, “freezing”, and “no-freezing” inhibition groups were tagged in the same way as their excitatory counterparts (see above).

### Inhibitory f-FLiCRE manipulation protocol

Optogenetic inhibition experiments were conducted in ctxA, one day after FC (i.e., during a fear memory recall session). Mice were placed in the chamber for a total of 8 minutes, receiving yellow light stimulation in alternating 2-minute bouts (2 minutes OFF, 2 minutes ON, 2 minutes OFF, 2 minutes ON, continuous).

### Experiment control

Whenever a closed-loop setup was not needed (e.g when tagging shocks, during optogenetic manipulation, etc.), we used Bonsai ^45^ to synchronize camera recording, light delivery, and shocks in the case of training sessions, via an Arduino Due running the Firmata protocol.

### Closed-loop optogenetics

For behavior-dependent tagging experiments (i.e., whenever tagging happened during freezing bouts or outside of freezing bouts), we used a combination of DeepLabStream ^18^ and custom Python code to track mice during the experiment with our pre-trained DeepLabCut ^46^ network. Each frame acquired live by the webcam was analyzed at 15 fps, and 10 body parts were tracked. Each of these body parts’ speed was computed as the distance the body part moved since the previous frame, divided by 1/15 seconds. For each frame, each body part was considered either mobile or immobile using a threshold of 0.5 cm/s. To ensure that the freezing tag groups contained as much actual freezing as possible, and conversely, for the no-freezing groups, we used two different thresholds of freezing: for the freezing groups, mice were considered freezing if 7 or more body parts were immobile. For the no-freezing groups, mice were considered freezing if 6 or more body parts were immobile. This labeling was then transmitted through an “Arduino Due” running the Firmata protocol to control the tagging laser accordingly. Both the raw video and live tracking were saved for subsequent analysis.

### Seizure tagging experiment

Seizure tag animals received an i.p. injection of 20mg/kg kainic acid – a glutamate agonist – to induce epileptic seizures as described previously. ^4^. Animals were placed in an isolated chamber and monitored for seizure levels using the mouse-modified Racine scale ^47^. After about 1 hour, animals reached stage 5-6 seizures (“rearing and falling with forelimb clonus”), and blue light was shone continuously for one minute, 5 separate times, for a total of 5 minutes of tagging over about 15 minutes. Animals were then allowed to recover under close monitoring (seizures decrease in intensity and stop after one to two hours) and sacrificed the next day by intracardiac perfusion (see below).

### Perfusions and immunohistochemistry

Mice were overdosed with Ketamine/ Xylazine (200/20 mg/kg) and perfused transcardially with 30 mL of 0.9% saline, followed by 30 mL of 4% paraformaldehyde (PFA) in saline. Brains were carefully extracted and placed in 4% PFA at 4°C for 24 hours. They were then transferred to 30% sucrose until they sank to the bottom (1-2 days). Brains were then cut with a cryostat in 50µm coronal slices. Slices were stained with DAPI (1:10000 in PBS for 5 minutes) before being mounted on microscope slides with PermaFluor (Thermofisher).

### Imaging

Brain slices were imaged with a confocal microscope (Nikon A1 or Leica SP8), using 3 lasers: 405 nm (DAPI), 488 nm (eGFP) and 561 nm (mCherry). Laser strength was manually adjusted in between slices and animals to match background levels of fluorescence on all 3 lasers.

### General f-FLiCRE cell counting

We imaged 4 to 6 slices per animal, focusing on slices with the best combination of optrode placement and infection quality. Animals without suitable slices (due to subpar fiber placement, poor infection, or both) were excluded from further analyses. To determine the proportion of tagged cells in each group, cells were detected using QuPath 0.4.0 ^48^’s positive cell detection function. Cell bodies were identified from the green channel (GFP-CaM-uTEVp+ cells), and positivity was assessed from the averaged red channel value within each detected cell body (mCherry+ cells). The same parameters were used for all slices, except for the threshold, which was manually adjusted for each slice to match the experimenter’s manual counting. Cell counts were then unblinded and pooled across animals.

### Post-hoc analysis of freezing

Top-down video recordings of behavior were analyzed using pre-trained DeepLabCut networks (ResNet152-based, 1000 training images, 1M training iterations) to obtain frame-by-frame tracking of 10 body parts (nose, neck, ears, sides, middle-back, hindlegs, tail base). This tracking data was then input into BehaviorDEPOT ^49^ for automatic freezing detection. The “freezing_velocity” classifier was used to identify freezing bouts in the videos, combining a threshold on overall animal speed and head movement speed with the following parameters:

● Velocity threshold: 0.3
● Angle threshold: 12
● Window width: 32
● Count threshold: 10
● Min. duration: 0.5s

This automatic freezing scoring was validated by comparing it to a blinded experimenter’s scoring of five behavior videos (Fig.S1C-D).

### In-vivo calcium imaging viral preparation

AAV9-packed syn-jGCaMP8f-WPRE was ordered from Addgene, using a plasmid gifted by the GENIE project (Addgene viral prep #162376-AAV9). The stock viral preparation was diluted 1:1 with saline before surgeries to achieve a titer of approximately 10^12^ viral particles per mL.

### In-vivo calcium imaging surgeries

For all surgeries, mice were anesthetized with an i.p. injection of Ketamine/Xylazine (100/10 mg/kg) and received local anesthesia via subcutaneous injection of lurocaïne (4 mg/kg). Analgesia was provided via subcutaneous injection of buprenorphine (0.1 mg/kg). After each surgery, mice were rehydrated and received further analgesia through subcutaneous injections of 5% glucose water (150µL) and Meloxicam (20 mg/kg).

### GCaMP injection and lens implantation

This surgery was performed following previously described methods ^50^. Briefly, a 1mm wide craniotomy was made at -2mm AP and -1.5mm ML. The dura was opened, and the cortex was aspirated until crossed fibers were observed (around -1.1DV). After aspiration, 1µL of AAV9-syn-jGCaMP8f-WPRE was injected into the dCA1 at -2mm AP, -1.3mm ML, -1.4mm DV. A GRIN lens (GoFoton 1mm lens) was then implanted at -2mm AP, -1.5mm ML, -1.3mm DV and secured with cement and screws.

### Baseplate implantation

The baseplate was implanted while attached to the miniscope to visualize the fluorescent field of view for correct positioning. Animals with discernible vasculature and activity had the baseplate secured to the existing lens cement with superglue and additional black cement. For animals with blurry fluorescence and vasculature, baseplating was attempted again one or two weeks later. If no fluorescence was observed, these animals were kept as companions and sacrificed at the same time as the experimental animals.

### In-vivo calcium imaging experimental timeline

On day 1, miniscope-implanted animals were first imaged in a neutral context (ctxB). On day 2, they underwent fear conditioning in ctxA (FC ctxA) and were tested on day 3 (RECALL ctxA). On day 4, they were imaged in another neutral context (ctxC).

### Lens placement and GCaMP infection validation

After finishing all the imaging sessions, mice were perfused, and brain slices were collected and imaged as previously described (i.e., DAPI-stained and imaged with a confocal microscope). For each animal, we verified that dCA1 cells were adequately infected (GCaMP fluorescence can be observed ex-vivo using a 488mm laser). GRIN lens placement was verified by identifying the hole it left in the brain and checking its alignment and distance to the granular layer (optimal placement is ∼100-200 µm above and aligned with the imaged cells).

### Post-hoc analysis of calcium traces

We visualized the activity of N=9 animals, with an average 90 cells per session (min=47, max=172, median=90, quartile1=64, quartile3=109). Calcium traces were extracted from saved videos with Min1pipe v3.1 ^51^ running in Matlab R2021b. Traces were manually inspected before analysis to remove false positives and duplicates.

### Detection of cell groups in FC

During FC, the analysis of calcium traces was matched to the f-FLiCRE experimental procedure. For any given group (e.g. “preshock”), calcium traces were averaged during the corresponding period (e.g. 2 minutes of preshock) that would have been the FLiCRE tagging period. Cells were then ranked according to this average, and the distribution obtained. All cells above the mean plus one standard deviation were designated as belonging to that particular group.

### RECALL cells

Similarly to the FC cell groups, the calcium activity of each cell during the recall session was averaged during the freezing periods (“Frz RECALL” group) or no-freezing periods (“No-Frz RECALL” group) of recall. Cells were ranked from highest to lowest, and cells above mean plus one standard deviation were selected.

### Cell population overlaps

We computed overlaps between these four cell populations. For statistical analysis, we compared these real overlaps to a bootstrapped distribution of overlaps (i.e., chance-level overlap). Cell populations were considered significantly overlapped if their real overlap fell above the 95th percentile of the corresponding bootstrapped distribution. Conversely, they were considered significantly non-overlapped if their real overlap fell below the 5th percentile of the corresponding bootstrapped distribution.

### Cell registration across experiments

Cells were tracked across different experiments using CellReg v1.5.5 ^52^, running in Matlab R2021b. Briefly, neuronal footprints from all experiments are obtained from the Min1pipe analysis. Experiments are then aligned using translations and rotations. The resulting shifted footprints were compared between sessions using spatial correlation, and a probabilistic model was fitted to these correlations to determine positive and negative matchings. We used the following parameters for all sessions:

● Alignment type: Translations and rotations
● Model maximal distance: 14 microns
● Initial registration type: Spatial correlation
● Initial threshold: 0.5
● Registration approach: Probabilistic modeling
● Final registration type: Spatial correlations
● P_same threshold: 0.5

### FC cells in recall

Cells tracked from FC ctxA to recall were sorted into their respective groups from FC: “pre-shock”, “shock”, “freezing”, “no-freezing”, and others (where others contained all cells detected in both FC and in recall that did not fall into the other four categories). For each animal, we calculated the relative number of cells detected during recall per FC group. This was computed as the percentage of cells from that FC group present in recall, divided by the overall percentage of FC cells detected in recall. For example, if mouse N has 100 neurons detected in FC and 70 are also detected in recall (70% overall percentage), and out of the 10 “pre-shock” cells detected in FC, 5 are detected in recall (50% “pre-shock” percentage), the relative number of cells would be: 50%/70% = 0.71. Hence, chance level corresponds to a relative value of 1.

### Binned activity

To compare activity between cells and compute correlations, calcium traces were binned in 1-second bins and z-scored to remove any activity below 2 SD. Peaks were then detected using the Matlab Findpeaks function to determine the position of calcium ‘events’, and create binned arrays of events (0 for a second without events, 1 for a second with at least one event).

### Overall cell activity

Overall cell activity was computed as the average number of calcium events per second, using the binned data. Specifically, we computed the average value of the binned array of events for each cell.

### Correlations

We obtained Pearson correlations of the binned calcium traces for every cell pair. Correlations can be compared between the groups by extracting from the full matrix of cell-pair correlations any cell-pair correlation belonging to group A-group B. For example, one can observe the “group intra-correlations” of freezing cells by only looking at correlations of freezing-freezing cell pairs. Furthermore, one can observe “group inter-correlations” by looking at correlations of freezing-shock cell pairs. These comparisons were done in FC, RECALL ctxA, and ctxC for the cell groups detected in FC (FC “pre-shock” cells, FC “shock” cells, FC “freezing” cells, FC “no-freezing” cells) either directly (in the FC) or by using CellReg to track the cells in the other experiments (RECALL ctxA and OF).

### Random correlations

Random correlations were computed by circularly shifting the event arrays by a random amount 1000 times and correlating these random event arrays together. Each group’s intra- and inter-correlations were then compared to the corresponding random correlations to control for sample size effects in the direct comparisons.

### Topological analysis

Following analysis of FC ctxA activity, neurons from the different groups were traced back to their position in the miniscope’s field of view during FC ctxA. To determine the average distance of neurons in group X to the closest neuron in group Y, the distance of each neuron in group X to every neuron in group Y was computed, and only the smallest of these distances was kept. These “closest neighbor distances” were then averaged per animal, and statistical analysis was done by comparing the average values per animal. Note that the average distance of the closest neighbor X->Y can differ from that of Y->X.

### Detection of cell groups - statistical approach

To validate miniscope results obtained with the “activity ranking method’, we applied all the same analyses to cells assigned to groups using a statistical criterion instead. For every group, for every cell, we obtain a random distribution of its calcium trace by applying 1000 random circular shifts to it. We then compare activity between the “in”, i.e. putative FLiCRE tagging for the corresponding group, and “out” period (everything else), for the real trace and all the random ones. A cell is designated as belonging to the corresponding group if the real trace’s in/out ratio of activity exceeds that of 95% of the associated random distribution. Subsequent analyses were as described above.

#### SUPPLEMENTARY MATERIAL

**Fig.S1.**
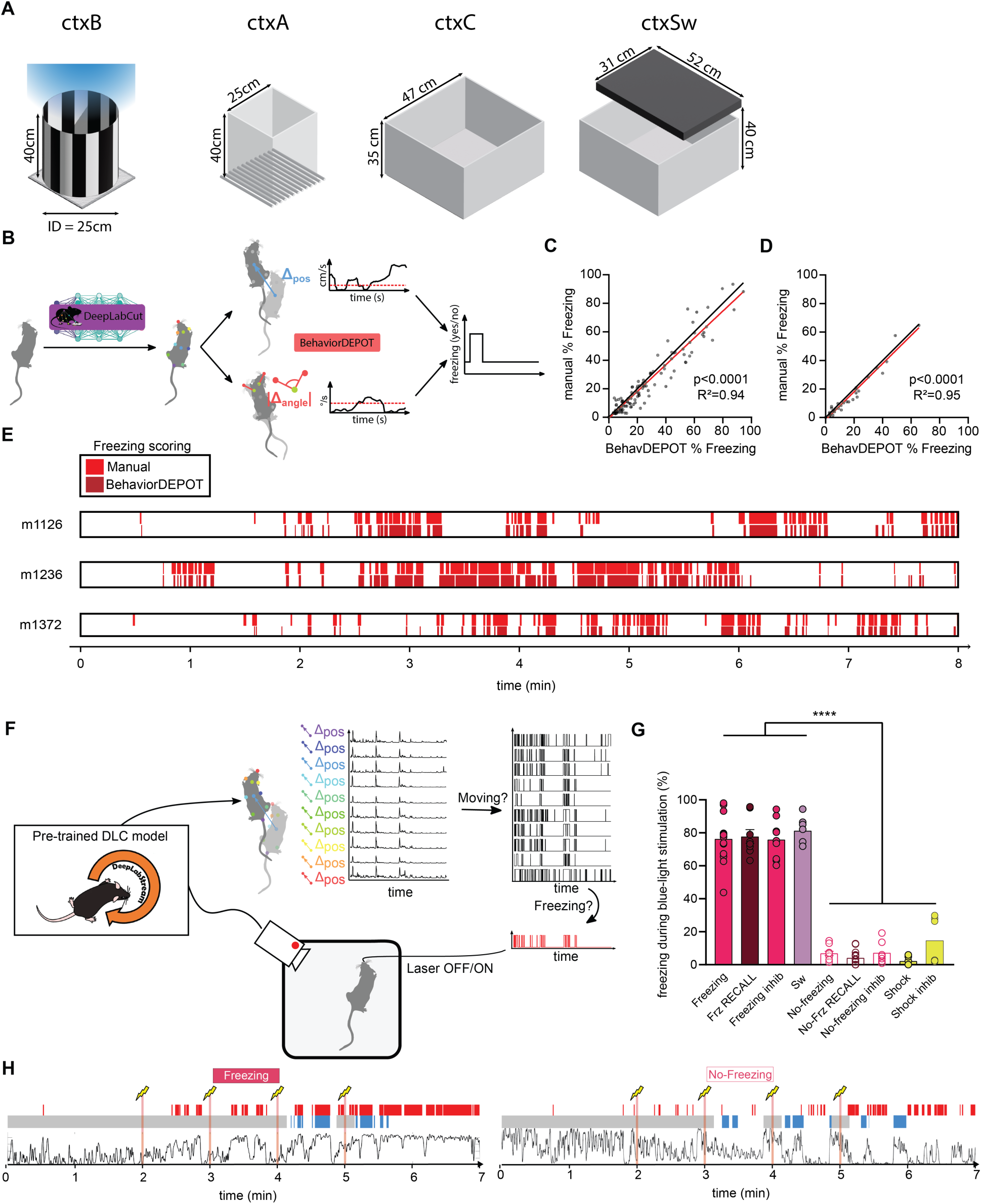
Offline and live automatic freezing scoring methods. (**A**) Schematics of the different contexts used. (**B**) DeepLabCut and BehaviorDEPOT pipeline for automatic freezing scoring. (**C**) Comparison of automatic and manual freezing scoring with videos split in 20s bouts or (**D**) in 2-minute bouts (simple linear regressions). (**E**) Example animal freezing scores during ctxC using manual versus behaviorDEPOT scoring. (**F**) DeepLabStream pipeline for freezing/no-freezing detection dependent closed-loop tagging. (**G**) Percentage of freezing during blue light tagging for the different groups. (**H**) Example of animals’ post-hoc freezing scoring and blue light tagging during FC ctxA. Each data point corresponds to the mean value for an individual animal while bars represent mean ± SEM across animals. ****p<0.0001.

**Fig.S2.**
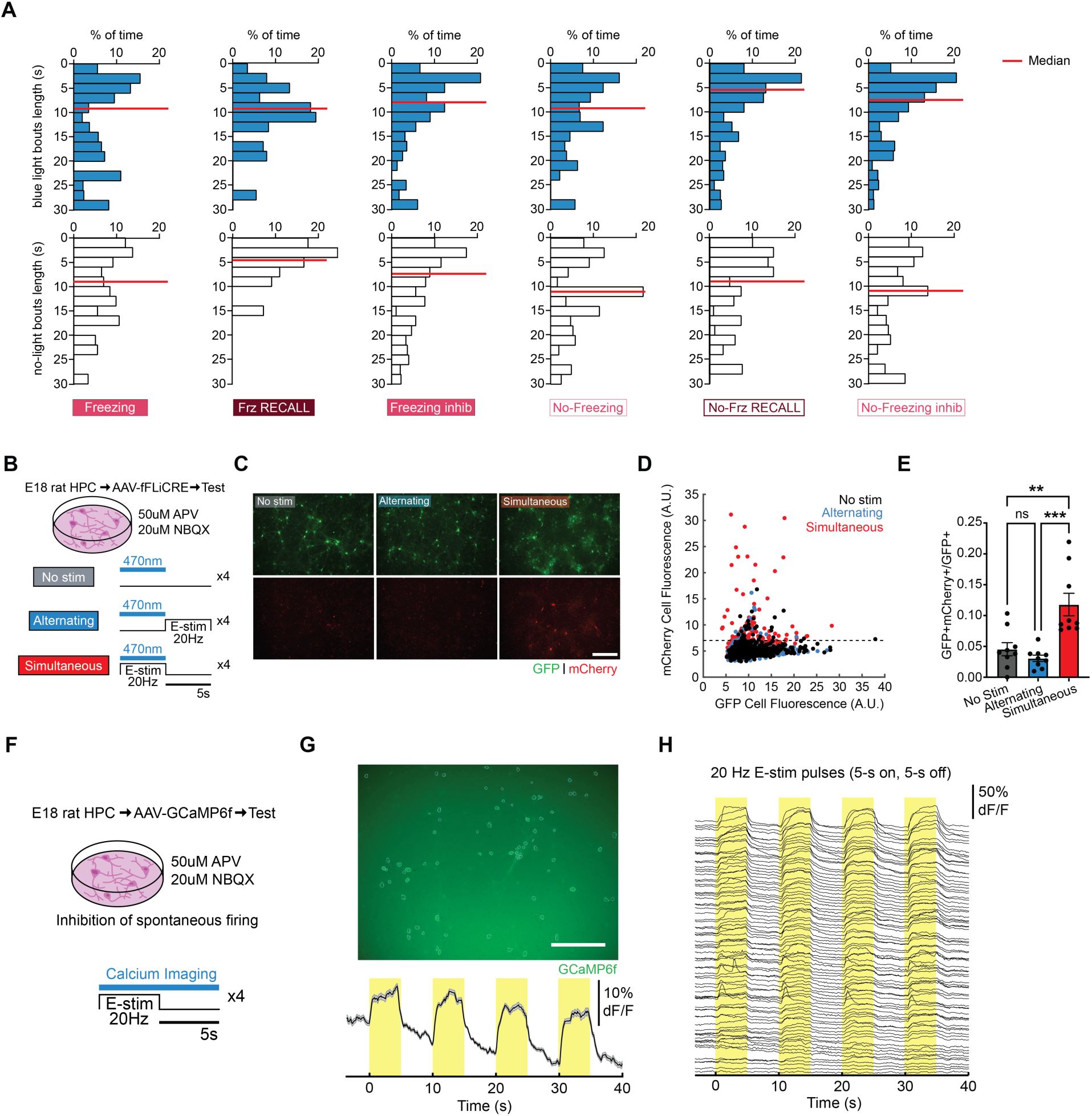
In-vitro validation of the novel f-FLiCRE tool. (**A**) Weighted histograms representing proportionality of blue-light bouts (blue) or no-light bouts (white) for freezing/no-freezing groups. Median in red. (**B**) Schematic of the protocol for in-vitro f-FLiCRE validation. Neurons expressing f-FLiCRE proteins have their spontaneous firing inhibited with 50 µM APV and 20 µM NBQX, and are stimulated with an electric field at 20Hz for 5 seconds at a time. (**C**) Example FOVs for all three experimental conditions. Scale bar: 100 µm. (**D**) mCherry vs. GFP cell fluorescence from segmented cells. Dotted line represents threshold for positive counting. (**E**) Fraction of GFP+mCherry+/GFP+ cells for the three conditions. (**F**) Schematic of the protocol for in-vitro stimulation/inhibition validation. (**G**) Top: FOV of cultured rat hippocampal neurons expressing AAV5-hSyn-GCaMP6f and treated with electric field stimulation for 5-s on, 5-s off, for a total duration of 40s (5-ms pulse width, 20 Hz pulse frequency). Neurons were treated with 50 µM APV and 20 µM NBQX. Scale bar, 100µm. Bottom: Mean dF/F traces of *n* = 78 neurons detected in the FOV. (**H**) Individual traces of the responses of all 78 neuron GCaMP6f to the electric field stimulation. Each data point corresponds to the mean value of an individual FOV, while bars represent mean ± SEM across FOVs. Statistical test is an ordinary one-way ANOVA and Tukey’s multiple comparisons test. **p<0.01, ***p<0.001.

**Fig.S3.**
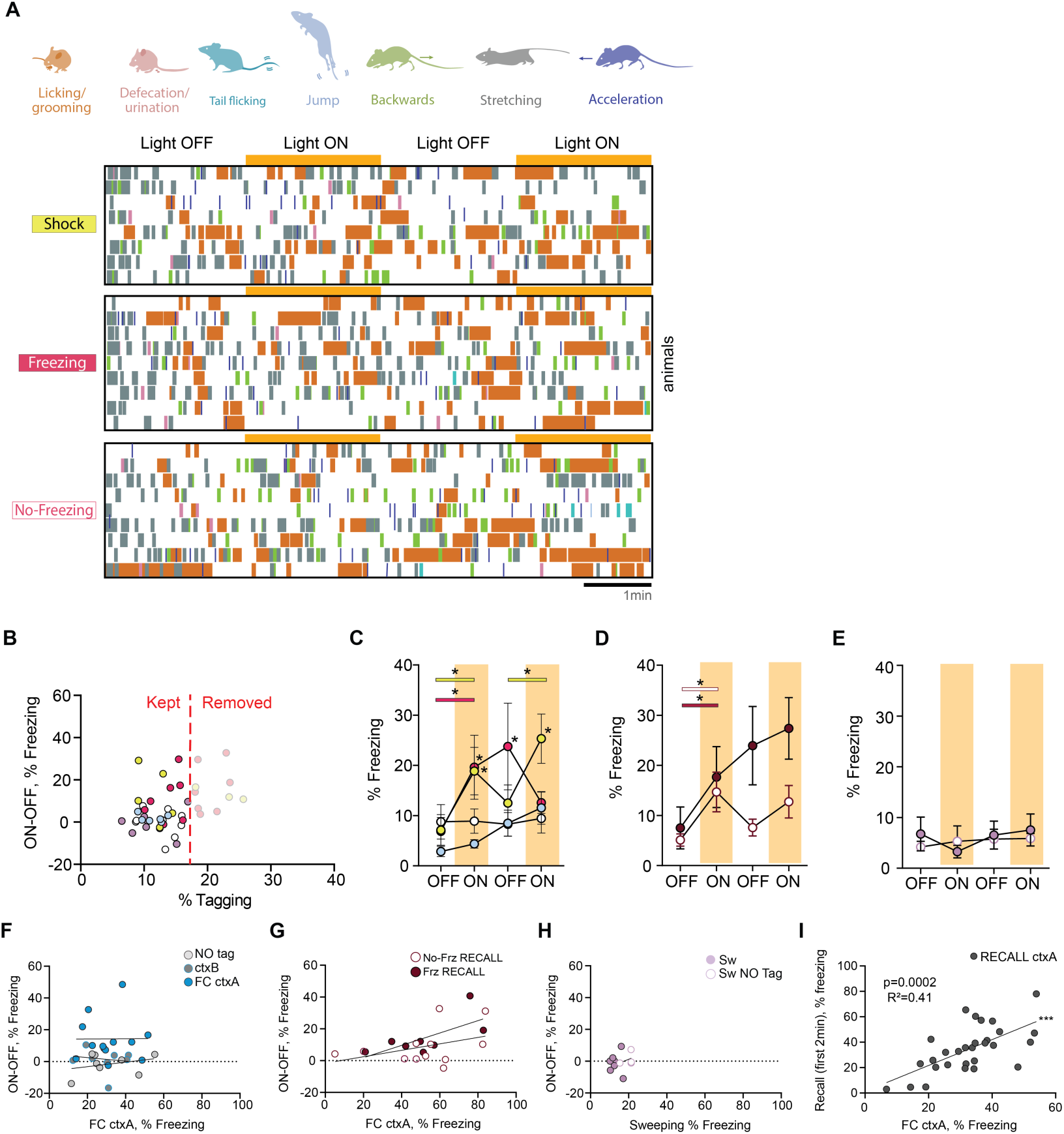
Additional behavioral tracking and counting. (**A**) Tracking of various behaviors during ctxC in some example groups and animals (RM one-way ANOVA for each type of behavior is non-significant). (**B**) Representation of cutoff % tagged value used to compare the behaviors of animals with similar % tagging values. Comparison of freezing levels between equivalently-tagged animals during light-OFF (no shading) and light-ON epochs (yellow shading) in (**C**) FC ctxA-tagged animals, (**D**) RECALL ctxA-tagged animals, and (**E**) sweeping-tagged animals. (**F**) Correlation between Δfreezing in ctxC and overall freezing in FC ctxA for Fig.1 groups and (**G**) Fig.2P-Q groups (simple linear regressions). (**H**) Correlation between Δfreezing in sweeping-test and overall freezing during sweeping for Fig.2R-S groups (simple linear regressions). (**I**) Correlation between freezing in the first 2 minutes of recall (i.e. before any optogenetic manipulation) and freezing in FC ctxA for all inhibitory groups. Statistical differences between groups are depicted with simple asterisks, while between epochs with color-coded lines, only for consecutive periods (non-consecutive significance not shown). Each data point corresponds to the mean value for each individual animal while bars represent mean ± SEM across animals. *p<0.05, ***p<0.001.

**Fig.S4.**
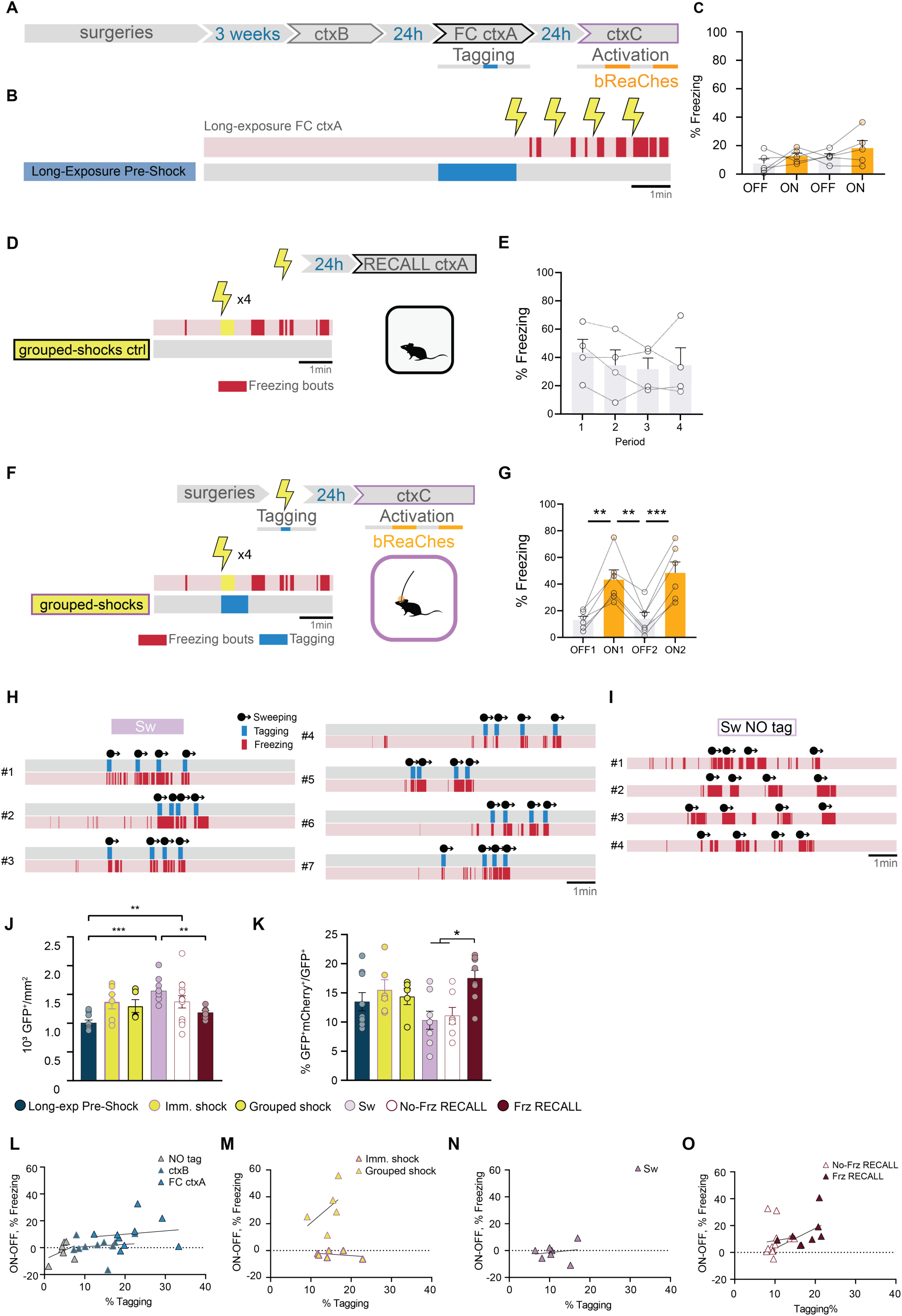
Long exposure, sweeping, and RECALL tagging experiments. (**A**) Long-exposure pre-shock experiment protocol and (**B**) tagging timeline. (**C**) Freezing levels of individual “long-exposure pre-shock” animals during ctxC, outside (gray) and during (yellow) light-on periods. (**D**) Schematic of grouped-shock ctrl experiment: After training in ctxA with the same shock patterns as immediate shock groups, freezing is tested in a recall ctxA session. (**E**) Freezing level in recall ctxA. (**F**) Schematic of grouped-shock experiment: after opto-tagging in ctxA with the same shock patterns as immediate shock groups, freezing is tested in an opto-reactivation test ctxC session.(**G**) Freezing level in test ctxC. (**H**) Timelines of all “Sw” and (**I**) “Sw NO tag” animals, displaying sweeping bouts, freezing, and tagging for the former. (**J**) Density of GFP+ (i.e. infected) neurons in the dCA1. (**K**) Percentage of tagged cells (GFP+mCherry+/GFP+). (**L**) Correlation between Δfreezing in ctxC/sweeping and % of tagged cells for Fig.1 groups. (**M**) Correlation between Δfreezing in ctxC and % of tagged cells for imm.shock and grouped shock groups (**N**) Correlation between Δfreezing in sweeping-test and % of tagged cells for “Sw” tagged animals. (**O**) Correlation between Δfreezing in ctxC/sweeping and % of tagged cells for Fig.2K1-L groups. Each data point corresponds to the mean value for each individual animal while bars represent mean ± SEM across animals. **p<0.01, ***p<0.001.

**Fig.S5.**
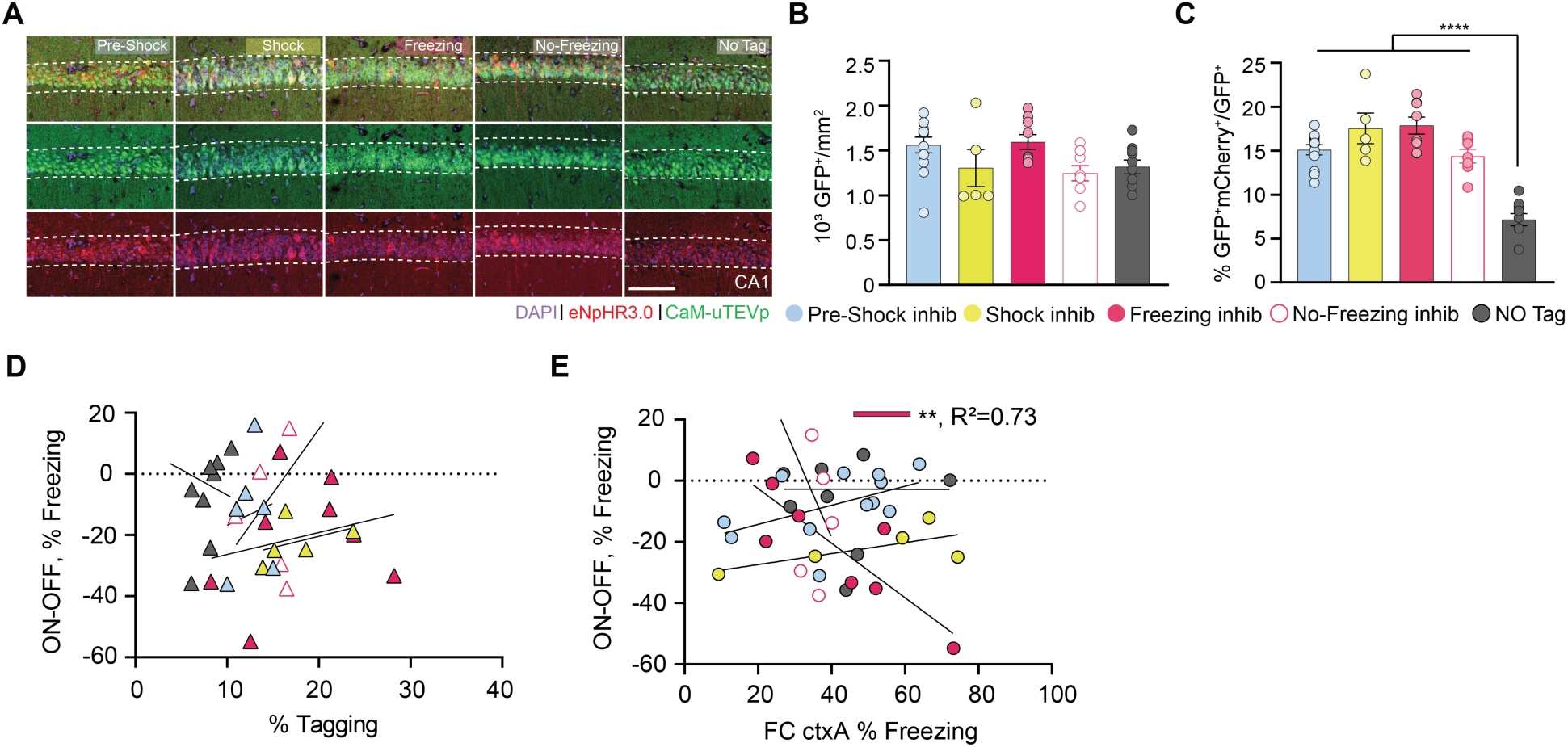
FLiCRE-tagging and behavior analysis in inhibitory groups. (**A**) Representative images of infection (green) and tagging (red) for inhibitory groups. Scale bar: 100µm. (**B**) Density of GFP+ (i.e., infected) neurons in the gCA1 for the inhibitory groups. (**C**) Percentage of tagged cells (GFP+mCherry+/GFP+). (**D**) Correlation between Δfreezing in RECALL ctxA and % of tagged cells for inhibitory groups. (**E**) Correlation between Δfreezing in RECALL ctxA and overall freezing in FC ctxA for inhibitory groups. Each data point corresponds to the mean value for an individual animal while bars represent mean ± SEM across animals. Statistical tests are ordinary one-way or linear regressions depending on context. **p<0.01, ****p<0.0001.

**Fig.S6.**
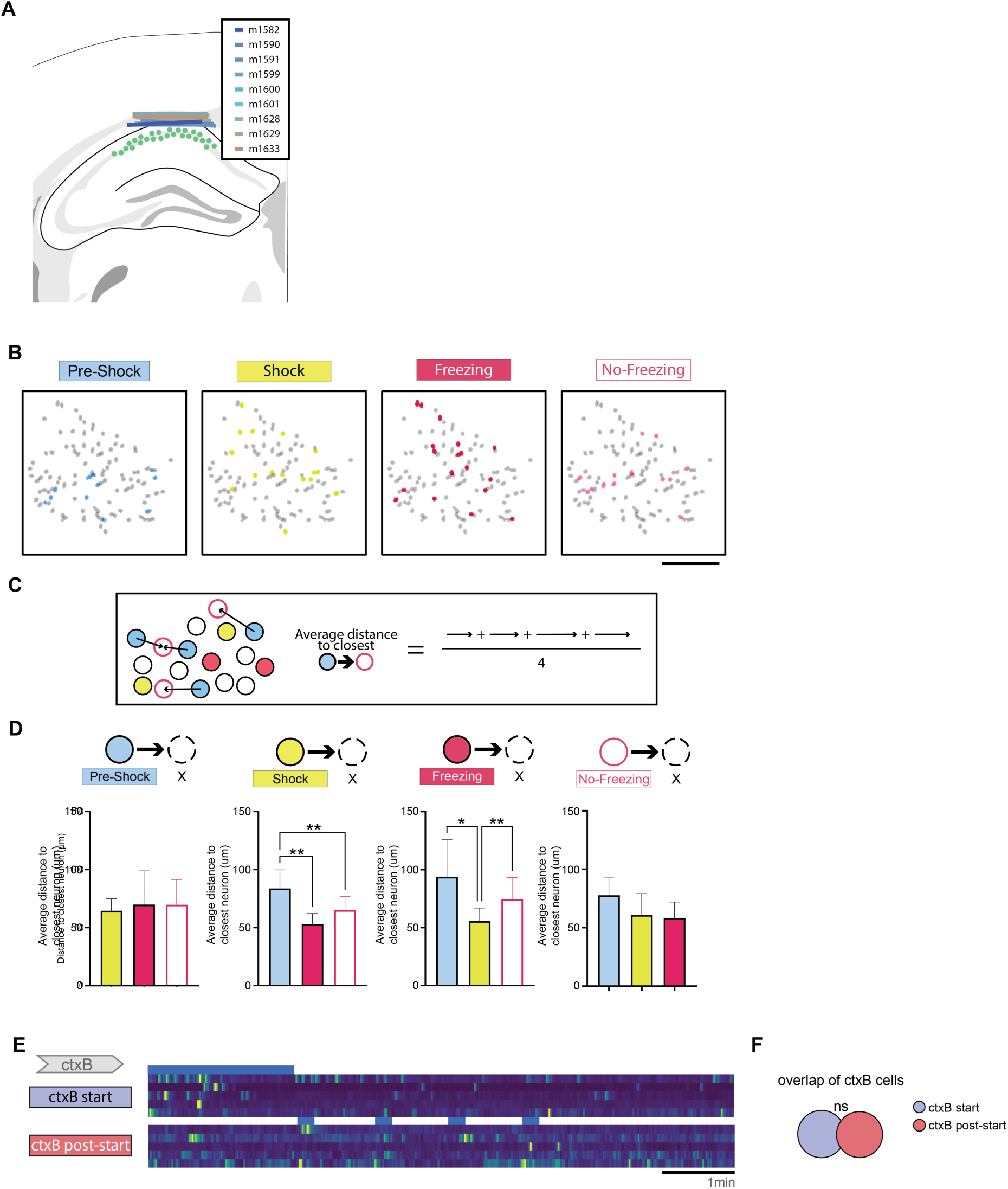
Calcium imaging: validation, overlap statistics, and topological analysis. (**A**) Schematic of lens positioning and infection for all 9 animals implanted for calcium imaging experiments. (**B**) Representative animal’s miniscope field of view, with the topology of the 4 tracked ‘FLiCRE-like’ groups shown. Scale bar: 200µm. (**C**) Computation of average distance to closest neighbor between two cell groups. (**D**) Average distance to closest neighbor between all pairs of cell groups (ordinary one-way ANOVA). (**E**) Example calcium traces from ctxB. Blue represents the periods used to determine which cells belong to the corresponding group, and would have been tagged using FLiCRE. (**F**) Overlap between cell groups of ctxB. Each data point corresponds to the mean value for an individual animal while bars represent mean ± SEM across animals. *p<0.05, **p<0.01.

**Fig.S7.**
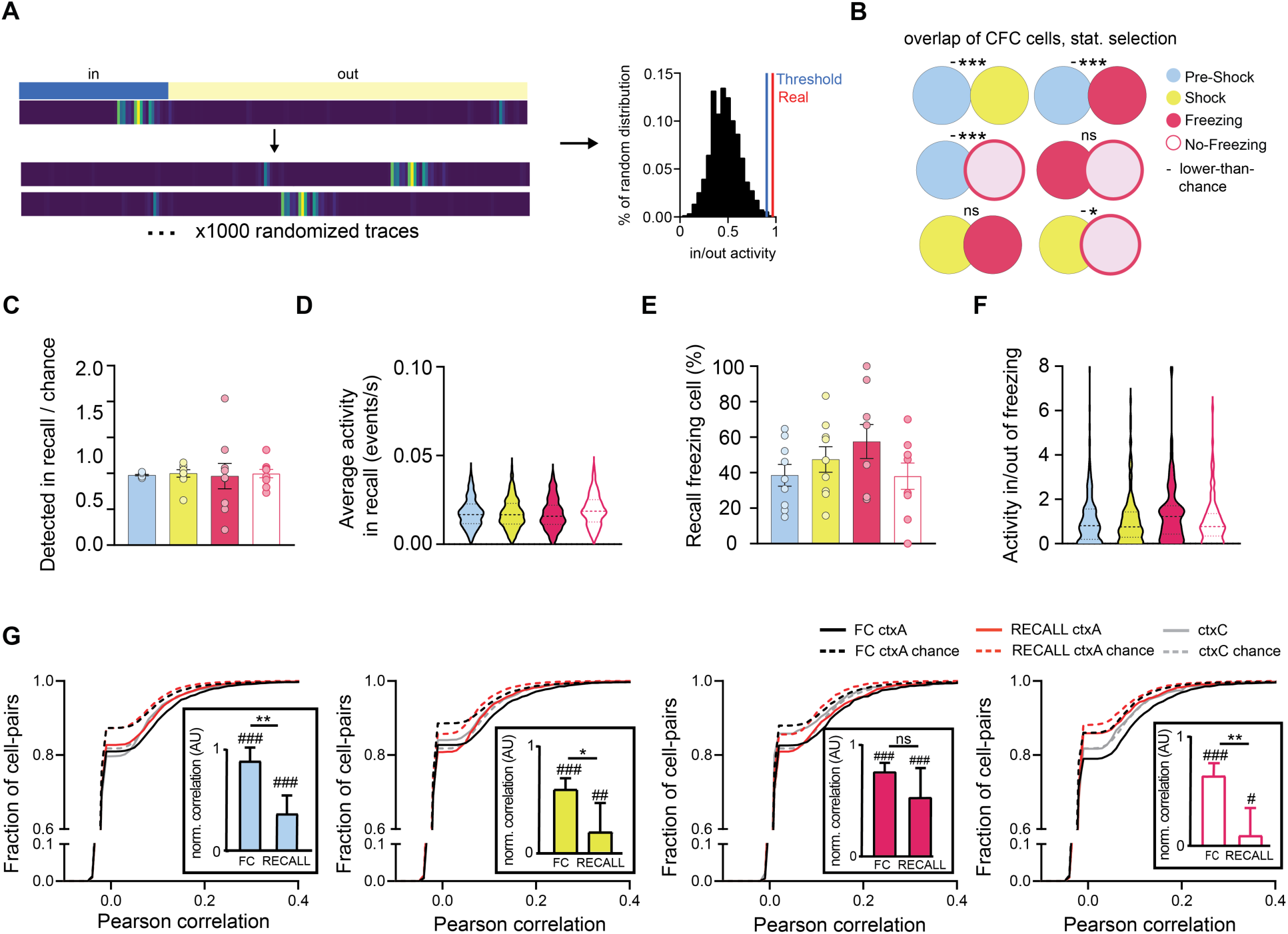
Calcium imaging: Analysis with statistical criteria. (**A**) Schematic of method for sorting cells into the “Pre-Shock”, “Shock”, “Freezing”, and “No-Freezing” groups using a statistical criterion (example shown for pre-shock). (**B**) Overlap between cell groups from the same experiment, top: FC, and bottom: recall. + denotes higher-than-chance overlaps, - lower than chance overlaps. (**C**) Number of cells per animal from FC ctxA cell groups detected in RECALL ctxA over chance number by cell group. (**D**) Average calcium events per second in RECALL ctxA for tracked cells by cell group. (**E**) Percentage of FC ctxA groups cells that are freezing cells in Test ctxA. (**F**) Activity ratios per animal of FC ctxA groups for cells inside/outside of Test ctxA freezing bouts. (**G**) Cumulative distributions of same-group cell-pair Pearson correlations in FC ctxA (black lines), RECALL ctxA (red lines) and ctxC (gray lines). Chance-level correlations are shown with dotted lines. Statistical tests are comparisons to bootstrapped distributions for statistical selection and for overlaps, and ordinary one-way ANOVA otherwise. *p<0.05, ***p<0.001, #p<0.05, ###p<0.001.

**Movie S1. Example ctxC session of “Pre-Shock” tagged animal.** Excerpt from test session in ctxC, from minute 1 to minute 3 (i.e. one minute light OFF, one minute with light ON), sped up 4 times. Freezing is indicated as scored by BehaviorDEPOT.

**Movie S2. Example ctxC session of “Shock” tagged animal.** Excerpt from test session in ctxC, from minute 1 to minute 3 (i.e. one minute light OFF, one minute with light ON), sped up 4 times. Freezing is indicated as scored by BehaviorDEPOT.

**Movie S3. Example ctxC session of “Freezing” tagged animal.** Excerpt from test session in ctxC, from minute 1 to minute 3 (i.e. one minute light OFF, one minute with light ON), sped up 4 times. Freezing is indicated as scored by BehaviorDEPOT.

**Movie S4. Example ctxC session of “No-Freezing” tagged animal.** Excerpt from test session in ctxC, from minute 1 to minute 3 (i.e. one minute light OFF, one minute with light ON), sped up 4 times. Freezing is indicated as scored by BehaviorDEPOT.

